# Dissecting and Tuning Primer Editing by Proofreading Polymerases

**DOI:** 10.1101/2021.05.11.443694

**Authors:** Daryl M. Gohl, Benjamin Auch, Amanda Certano, Brice LeFrançois, Anne Bouevitch, Evgueni Doukhanine, Christina Fragel, Jean Macklaim, Emily Hollister, John Garbe, Kenneth B. Beckman

## Abstract

Proofreading polymerases have 3’ to 5’ exonuclease activity that allows the excision and correction of mis-incorporated bases during DNA replication. In a previous study, we demonstrated that in addition to correcting substitution errors and lowering the error rate of DNA amplification, proofreading polymerases can also edit PCR primers to match template sequences. Primer editing is a feature that can be advantageous in certain experimental contexts, such as amplicon-based microbiome profiling. Here we develop a set of synthetic DNA standards to report on primer editing activity and use these standards to dissect this phenomenon. The primer editing standards allow next-generation sequencing-based enzymological measurements, reveal the extent of editing, and allow the comparison of different polymerases and cycling conditions. We demonstrate that proofreading polymerases edit PCR primers in a concentration-dependent manner, and we examine whether primer editing exhibits any sequence specificity. In addition, we use these standards to show that primer editing is tunable through the incorporation of phosphorothioate linkages. Finally, we demonstrate the ability of primer editing to robustly rescue the drop-out of taxa with 16S rRNA gene-targeting primer mismatches using mock communities and human skin microbiome samples.

## INTRODUCTION

Starting with the discovery of Taq polymerase (1) and the advent of the Polymerase Chain Reaction (PCR) (2, 3), lowering the error rate and improving the efficiency of DNA amplification have been ongoing experimental goals (4). These goals have been achieved through the discovery of additional naturally occurring polymerases with desirable properties, such as proofreading activity, as well as through engineering polymerases with desirable properties such as improved fidelity, specificity, efficiency, thermostability, inhibitor tolerance, reduced GC bias, and higher processivity (5).

Proofreading polymerases have 3’ to 5’ exonuclease activity, which allows the detection, excision, and replacement of mis-incorporated bases (6, 7). Such proofreading activity serves to minimize the rate of PCR errors and modern engineered enzymes have reported substitution error rates as low as 10^−6^-10^−7^ (8). We previously found that in addition to reducing the rate of amplification errors during PCR, proofreading polymerases are able to edit primer sequences to correct mismatches between the primer and template during amplification (9).

Primer editing activity can be used to prevent the dropout of taxa due to mismatches between the amplification primers and the DNA template in amplicon-based microbiome profiling. In such experiments, PCR primers are designed that target highly conserved regions of the 16S ribosomal RNA (rRNA) gene, the 18S rRNA gene, the Internal Transcribed Spacer (ITS) region, the mitochondrial Cytochrome Oxidase I (COI) gene, or other marker genes of interest. However, the binding sites for these “universal” primers are not perfectly conserved and even with the incorporation of degenerate bases, mismatches between the amplification primer and bacterial template sequences particularly in the critical last 3-4 bases of the primer can dramatically reduce amplification efficiency or even cause taxa to be undetected (10, 11). *In silico* analyses have found that for a set of 8 commonly used 16S rRNA gene-targeting primers, rates of primers having a mismatch in the last 4 nucleotides were greater than 10% in the majority of shotgun metagenomic datasets analyzed (12). In addition, the number of microbial taxa identified through shotgun metagenomic sequencing continue to grow (13, 14), leading to the identification of new taxa with divergent 16S rRNA gene sequences (15). Thus, methods to improve the recovery of taxa in spite of mismatches in marker gene amplification primers will help improve the accuracy of amplicon-based microbiome profiling.

In this study, we develop a set of synthetic DNA standards to report on primer editing activity and use them to characterize this phenomenon in more detail. Using this next-generation sequencing-based enzymology approach, we find that many proofreading polymerases are able to mediate primer editing, though to different extents, that primer editing can be tuned by adjusting enzyme concentration or through the incorporation of phosphorothioate bonds, and we demonstrate that primer editing can robustly and quantitatively rescue the dropout of key taxa in 16S rRNA gene microbiome profiles from human skin microbiome samples.

## MATERIAL AND METHODS

### Design, synthesis and cloning of primer editing standards

Primer editing standards were designed consisting of 332 bp of the *E. coli* 16S rRNA gene spanning variable region 4, the V4_515F and V4_806R primer sites, and 20 bp of flanking sequence. 31 different constructs were designed, with either the wild-type *E. coli* V4_515F primer binding site or each possible single base mismatch within the last 10 bp of the *E. coli* V4_515F primer binding site (Figure 1). In addition, the constructs contain an I-*Sce*I site for plasmid linearization, as well as a REcount PCR-free quantification barcode (16) for accurately measuring the relative construct abundance within the standard pool. See Supplemental File 1 for primer editing construct designs. Constructs were synthesized as DNA tiles by SGI-DNA and assembled and inserted into a pUCGA_1.0 backbone using the BioXP instrument. Successful assembly was verified by gel electrophoresis and plasmids were transformed into *E. coli* 5-alpha cells (NEB). Plasmid DNA from multiple independent clones was isolated for each construct using a Zymo Zyppy 96-well plasmid prep kit. Clones were sequenced by Sanger sequencing using the following sequencing primers and mutation-free clones for each construct were selected for further experiments: pUCGA1.0-Sanger_For: CGACTCTAGAGGATCGAGCACA pUCGA1.0-Sanger_Rev: TTCGAGCTCGGTACCCGCAT Sequence-verified clones were consolidated onto a single 96-well plate, normalized to 10 ng/ul with nuclease-free water, and pooled evenly (4 µl of each construct). The primer editing standard pool was diluted to a working concentration of 1 million molecules/construct/µl by diluting 5 µl of the 10 ng/µl standard pool in 452.95 µl of nuclease-free water.

**Figure 1.**
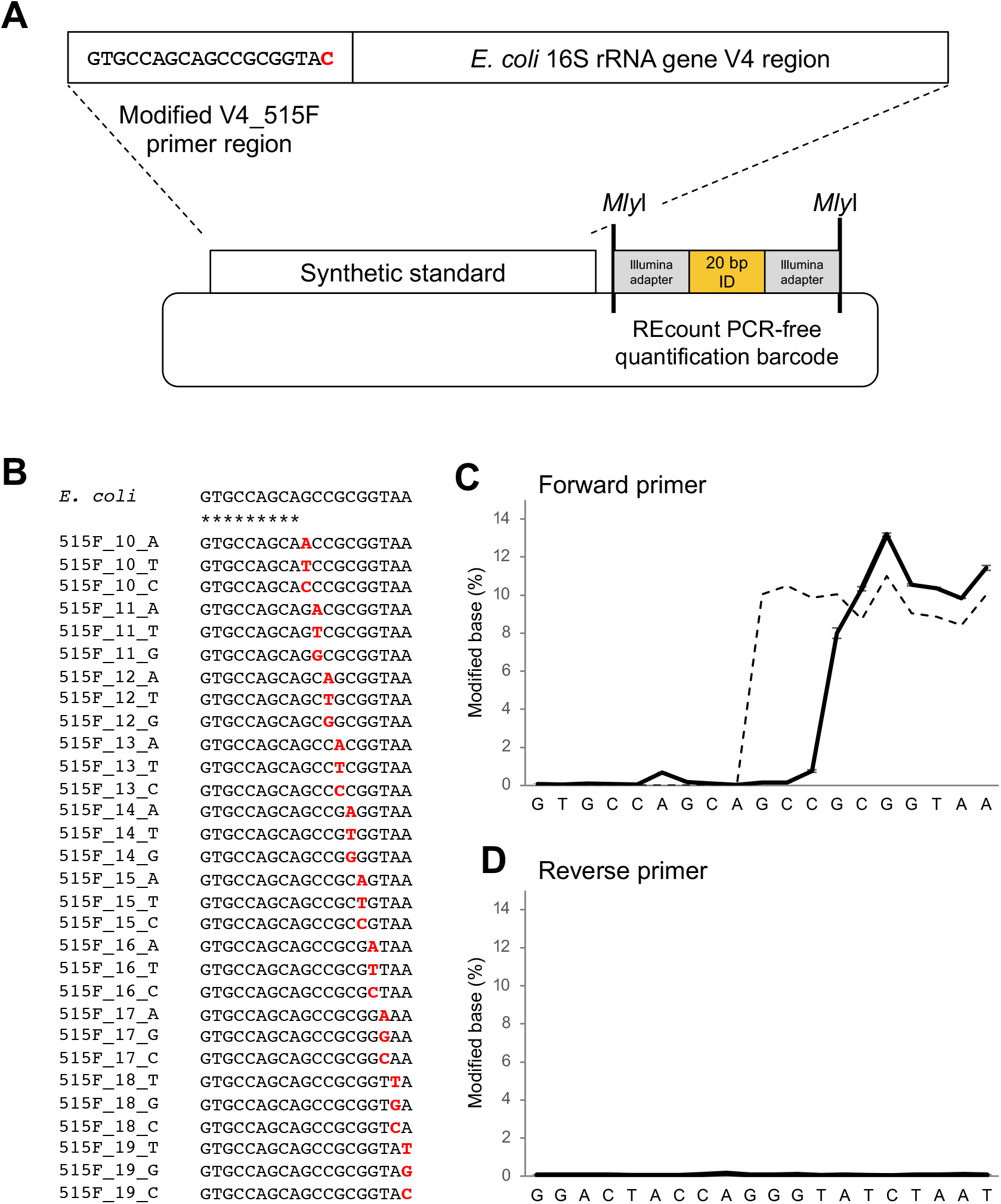
Synthetic DNA standards for measuring primer editing activity. A) Design of primer editing standard constructs. Each standard plasmid contains a copy of the *E. coli* 16S rRNA gene V4 region with a specific modification to the V4_515F primer binding region, as well as a REcount PCR-free barcode quantification construct for quantifying the abundance of each standard plasmid in the pooled mixture. B) The synthetic standard pool contains every possible single base substitution in the last 10 bp of the V4_515F primer binding region. C) Measured abundance of edited forward primer sequences when amplified with an *E. coli*-specific 515F/806R primer set and KAPA HiFi polymerase (solid line). Abundance of standard plasmids in the standard pool as assessed by REcount (dashed line). D) Measured abundance of edited primer reverse sequences when amplified with an *E. coli*-specific 515F/806R primer set.

### REcount-based quantification of primer editing standard pool

The following restriction digest was used to liberate the REcount PCR-free quantification barcodes: 17 µl primer editing standard pool DNA (10 ng/µl), 2 µl NEB Cutsmart buffer, 1 µl *Mly*I (NEB). The digest was incubated at 37**°**C for 1 hour. Added 30 µl of water to the digest (to bring the volume up to 50 µl). Added 30 µl of AMPureXP (Beckman Coulter) beads (0.6x), collected beads on a magnetic stand and transferred supernatant to new tube (discarded beads). Purified the supernatant using 1.8x AMPureXP beads (Beckman Coulter) and eluted in 25 µl of EB (Qiagen). The resulting sample was quantified using a Quant-iT PicoGreen dsDNA assay (Thermo Fisher Scientific), fragment sizes were assessed using an Agilent Bioanalyzer High Sensitivity assay and normalized to 2 nM for sequencing. The library was denatured with NaOH and prepared for sequencing according to the protocols described in the Illumina MiSeq Denature and Dilute Libraries Guide and sequenced in a portion of a MiSeq 600 cycle v3 run. Demultiplexed fastq files were generated using Illumina’s bcl2fastq software. REcount data was analyzed as previously described (16) using custom R and Python scripts and BioPython (17). The first 20 bp of the sequencing reads was mapped against a barcode reference file (Supplemental File 2), with a maximum of 2 mismatches allowed, using an analysis script which is available on GitHub (https://github.com/darylgohl/REcount).

### Testing different polymerases and polymerase concentrations

8 different polymerases (KAPA HiFi HotStart, Qiagen Taq, Q5 (NEB), Phusion Hot Start Flex DNA Polymerase (NEB), Vent (NEB), PfuUltra II Fusion HotStart DNA Polymerase (Agilent), Accuprime Taq (Thermo Fisher Scientific), and NEB Taq) were tested at 4 different concentrations (0.25x, 0.5x, 1x, and 2x manufacturer’s recommended concentration), with the primer editing standard pool at 4 different template concentrations (250,000, 25,000, 2,500, and 250 template molecules per construct). Nextera-tailed *E. coli* specific V4_515F and V4_806R primers (with no degenerate bases) were used to amplify the primer editing standards (16S-specific sequences in bold): E_coli_V4_515F: TCGTCGGCAGCGTCAGATGTGTATAAGAGACAG**GTGCCAGCAGCCGCGGTAA** E_coli_V4_806R: GTCTCGTGGGCTCGGAGATGTGTATAAGAGACAG**GGACTACCAGGGTATCTAAT** The following PCR recipes were used:

KAPA HiFi: 2.5 µl DNA template, 3.5 µl nuclease-free water, 2 µl 5x KAPA HiFi buffer (Kapa Biosystems), 0.3 µl 10 mM dNTPs (Kapa Biosystems), 0.5 µl DMSO (Fisher Scientific), 0.2 µl KAPA HiFi Polymerase (1x polymerase condition, Kapa Biosystems), 0.5 µl forward primer (10 µM, IDT), 0.5 µl reverse primer (10 µM, IDT). For the KAPA HiFi enzyme concentration tests the amount of enzyme added was adjusted appropriately (0.05 µl was added for the 0.25x condition, 0.1 µl was added for the 0.5x condition, and 0.4 µl was added for the 2x condition) and the amount of nuclease-free water was adjusted to compensate for the missing or added volume.

Q5: 2.5 µl DNA template, 3.7 µl nuclease-free water, 2 µl 5x Q5 buffer (NEB), 0.2 µl 10 mM dNTPs (NEB), 0.5 µl DMSO (Fisher Scientific), 0.1 µl Q5 Polymerase (1x polymerase condition, NEB), 0.5 µl forward primer (10 µM, IDT), 0.5 µl reverse primer (10 µM, IDT). For the Q5 enzyme concentration tests the amount of enzyme added was adjusted appropriately (0.025 µl was added for the 0.25x condition, 0.05 µl was added for the 0.5x condition, and 0.2 µl was added for the 2x condition) and the amount of nuclease-free water was adjusted to compensate for the missing or added volume.

Phusion: 2.5 µl DNA template, 3.7 µl nuclease-free water, 2 µl 5x Phusion buffer (NEB), 0.2 µl 10 mM dNTPs (NEB), 0.5 µl DMSO (Fisher Scientific), 0.1 µl Phusion Polymerase (1x polymerase condition, NEB), 0.5 µl forward primer (10 µM, IDT), 0.5 µl reverse primer (10 µM, IDT). For the Phusion enzyme concentration tests the amount of enzyme added was adjusted appropriately (0.025 µl was added for the 0.25x condition, 0.05 µl was added for the 0.5x condition, and 0.2 µl was added for the 2x condition) and the amount of nuclease-free water was adjusted to compensate for the missing or added volume.

Vent: 2.5 µl DNA template, 4.7 µl nuclease-free water, 1 µl 10x Vent buffer (NEB), 0.2 µl 10 mM dNTPs (NEB), 0.5 µl DMSO (Fisher Scientific), 0.1 µl Vent Polymerase (1x polymerase condition, NEB), 0.5 µl forward primer (10 µM, IDT), 0.5 µl reverse primer (10 µM, IDT). For the Vent enzyme concentration tests the amount of enzyme added was adjusted appropriately (0.025 µl was added for the 0.25x condition, 0.05 µl was added for the 0.5x condition, and 0.2 µl was added for the 2x condition) and the amount of nuclease-free water was adjusted to compensate for the missing or added volume.

Pfu: 2.5 µl DNA template, 4.7 µl nuclease-free water, 1 µl 10x Pfu buffer (Agilent), 0.1 µl 10 mM dNTPs (NEB), 0.5 µl DMSO (Fisher Scientific), 0.2 µl Pfu Polymerase (1x polymerase condition, Agilent), 0.5 µl forward primer (10 µM, IDT), 0.5 µl reverse primer (10 µM, IDT). For the Pfu enzyme concentration tests the amount of enzyme added was adjusted appropriately (0.05 µl was added for the 0.25x condition, 0.1 µl was added for the 0.5x condition, and 0.4 µl was added for the 2x condition) and the amount of nuclease-free water was adjusted to compensate for the missing or added volume. Accuprime Taq: 2.5 µl DNA template, 4.55 µl nuclease-free water, 1 µl 10x Accuprime buffer (Thermo Fisher Scientific), 0.2 µl 10 mM dNTPs (NEB), 0.5 µl DMSO (Fisher Scientific), 0.25 µl Accuprime Polymerase (1x polymerase condition, Thermo Fisher Scientific), 0.5 µl forward primer (10 µM, IDT), 0.5 µl reverse primer (10 µM, IDT). For the Accuprime Taq enzyme concentration tests the amount of enzyme added was adjusted appropriately (0.0625 µl was added for the 0.25x condition, 0.125 µl was added for the 0.5x condition, and 0.2 µl was added for the 2x condition) and the amount of nuclease-free water was adjusted to compensate for the missing or added volume.

NEB Taq: 2.5 µl DNA template, 4.75 µl nuclease-free water, 1 µl 10x Taq buffer (NEB), 0.2 µl 10 mM dNTPs (NEB), 0.5 µl DMSO (Fisher Scientific), 0.05 µl Taq Polymerase (1x polymerase condition, NEB), 0.5 µl forward primer (10 µM, IDT), 0.5 µl reverse primer (10 µM, IDT). For the NEB Taq enzyme concentration tests the amount of enzyme added was adjusted appropriately (0.0125 µl was added for the 0.25x condition, 0.025 µl was added for the 0.5x condition, and 0.1 µl was added for the 2x condition) and the amount of nuclease-free water was adjusted to compensate for the missing or added volume.

Qiagen Taq: 2.5 µl DNA template, 4.47 µl nuclease-free water, 1 µl 10x Taq buffer (Qiagen), 0.08 µl 10 mM dNTPs (NEB), 0.5 µl DMSO (Fisher Scientific), 0.4 µl MgCl2, 0.05 µl Taq Polymerase (1x polymerase condition, Qiagen), 0.5 µl forward primer (10 µM, IDT), 0.5 µl reverse primer (10 µM, IDT). For the Qiagen Taq enzyme concentration tests the amount of enzyme added was adjusted appropriately (0.0125 µl was added for the 0.25x condition, 0.025 µl was added for the 0.5x condition, and 0.1 µl was added for the 2x condition) and the amount of nuclease-free water was adjusted to compensate for the missing or added volume.

The following PCR cycling conditions were used:

KAPA HiFi: 95**°**C for 5 minutes, followed by 30 cycles of 98**°**C for 20 seconds, 55**°**C for 15 seconds, 72**°**C for 1 minute, followed by 72**°**C for 10 minutes.

Q5: 98**°**C for 30 seconds, followed by 30 cycles of 98**°**C for 20 seconds, 55**°**C for 15 seconds, 72**°**C for 1 minute, followed by 72**°**C for 5 minutes.

Phusion: 98**°**C for 30 seconds, followed by 30 cycles of 98**°**C for 20 seconds, 55**°**C for 15 seconds, 72**°**C for 1 minute, followed by 72**°**C for 5 minutes.

Vent: 95**°**C for 2 minutes, followed by 30 cycles of 95**°**C for 20 seconds, 55**°**C for 15 seconds, 72**°**C for 1 minute, followed by 72**°**C for 5 minutes.

Pfu: 95**°**C for 2 minutes, followed by 30 cycles of 95**°**C for 20 seconds, 55**°**C for 15 seconds, 72**°**C for 1 minute, followed by 72**°**C for 5 minutes.

Accuprime Taq: 95**°**C for 2 minutes, followed by 30 cycles of 95**°**C for 20 seconds, 55**°**C for 15 seconds, 68**°**C for 1 minute, followed by 68**°**C for 5 minutes.

NEB Taq: 95**°**C for 2 minutes, followed by 30 cycles of 95**°**C for 20 seconds, 55**°**C for 15 seconds, 68**°**C for 1 minute, followed by 68**°**C for 5 minutes.

Qiagen Taq: 95**°**C for 5 minutes, followed by 30 cycles of 94**°**C for 30 seconds, 55**°**C for 30 seconds, 72**°**C for 1 minute, followed by 72**°**C for 10 minutes.

Primary PCRs were then diluted 1:100 in sterile, nuclease-free water, and a second PCR reaction was set up to add the Illumina flow cell adapters and indices. All samples were indexed using KAPA HiFi polymerase using the following KAPA HiFi indexing PCR recipe: 5 µl 1:100 DNA template, 5 µl template DNA, 1 µl nuclease-free water, 2 µl 5x KAPA HiFi buffer (Kapa Biosystems), 0.3 µl 10 mM dNTPs (Kapa Biosystems), 0.5 µl DMSO (Fisher Scientific, Waltham, MA), 0.1 µl KAPA HiFi Polymerase (Kapa Biosystems), 0.5 µl forward primer (10 µM), 0.5 µl reverse primer (10 µM). Cycling conditions were: 95**°**C for 5 minutes, followed by 10 cycles of 98**°**C for 20 seconds, 55**°**C for 15 seconds, 72**°**C for 1 minute, followed by a final extension at 72**°**C for 10 minutes. The following indexing primers were used (X indicates the positions of the 8 bp indices): Forward indexing primer: AATGATACGGCGACCACCGAGATCTACACXXXXXXXXTCGTCGGCAGCGTC Reverse indexing primer: CAAGCAGAAGACGGCATACGAGATXXXXXXXXGTCTCGTGGGCTCGG The indexing PCR reactions were then purified and normalized using a SequalPrep normalization plate (Thermo Fisher Scientific), followed by elution in 20 µl of elution buffer. An even volume of the normalized libraries was pooled and concentrated using 1x AmpureXP beads (Beckman Coulter). Pooled libraries were quantified using a Quant-iT PicoGreen dsDNA assay (Thermo Fisher Scientific), fragment sizes were assessed using an Agilent Bioanalyzer High Sensitivity assay, and libraries were normalized to 2 nM for sequencing. The library was denatured with NaOH and prepared for sequencing according to the protocols described in the Illumina MiSeq Denature and Dilute Libraries Guide and sequenced in a portion of a MiSeq 600 cycle v3 run.

### Testing effects of annealing temperature

To test the effect of annealing temperature on primer editing, reactions were set up in triplicate for each of the 8 temperature conditions tested with KAPA HiFi HotStart or Qiagen Taq at 1x enzyme concentration using the recipes above. The primer editing standard pool was used at a template concentration of 10,000 template molecules/construct/µl (25,000 template molecules per construct in the amplification reaction). Reactions were run using the above cycling conditions for either KAPA HiFi or Qiagen Taq but using a gradient of annealing temperatures from 50**°**C to 60**°**C. The 8 temperatures tested were 50**°**C, 50.7**°**C, 52.1**°**C, 54**°**C, 56.2**°**C, 58.1**°**C, 59.4**°**C, and 60**°**C. The resulting amplicons were indexed using KAPA HiFi, normalized, quantified, and sequenced as described above.

### Amplification of wild type *E. coli* template with variant primers

A set of Nextera-tailed primers containing all 31 variants corresponding to those in the primer editing standards were designed (Supplemental File 3). These forward primers were pooled evenly and used to amplify a plasmid containing a tagged *E. coli* 16S rRNA template with wild-type primer binding sites, together with the E_coli_V4_806R reverse primer with either KAPA HiFi polymerase or Qiagen Taq polymerase across a gradient of annealing temperatures using the PCR recipes and cycling conditions described above. The resulting amplicons were indexed using KAPA HiFi, normalized, quantified, and sequenced as described above.

### Amplification with phosphorothioate-modified primers

To test the phosphorothioate primers, the primer editing standards were amplified at a template concentration of 10,000 template molecules/construct/µl (25,000 template molecules per construct in the amplification reaction) with either KAPA HiFi, Q5, or Phusion polymerase, at the 1x enzyme concentration using the recipes and cycling conditions above. The following exonuclease-protected primers were used, together with the the E_coli_V4_806R reverse primer (16S-specific sequences in bold, “*” indicates the position of a phosphorothioate bond):

E_coli_V4_515F_*19: TCGTCGGCAGCGTCAGATGTGTATAAGAGACAG**GTGCCAGCAGCCGCGGTA*A**

E_coli_V4_515F_*18: TCGTCGGCAGCGTCAGATGTGTATAAGAGACAG**GTGCCAGCAGCCGCGGT*AA**

E_coli_V4_515F_*17: TCGTCGGCAGCGTCAGATGTGTATAAGAGACAG**GTGCCAGCAGCCGCGG*TAA**

E_coli_V4_515F_*16: TCGTCGGCAGCGTCAGATGTGTATAAGAGACAG**GTGCCAGCAGCCGCG*GTAA**

E_coli_V4_515F_*15: TCGTCGGCAGCGTCAGATGTGTATAAGAGACAG**GTGCCAGCAGCCGC*GGTAA**

The resulting amplicons were indexed using KAPA HiFi, normalized, quantified, and sequenced as described above.

### Amplification of HM-276D mock community using V1V3, V4, V3V4, and V4V6 primers

The HM-276D mock community DNA was obtained through BEI Resources, NIAID, NIH: Genomic Mock Community B (HM-276D, Even, High Concentration, v5.1H). Reactions were set up in triplicate for each of the 4 primer sets with KAPA HiFi HotStart or Qiagen Taq at 1x enzyme concentration using the recipes and cycling conditions described above, with the following differences:

For V1V3 amplification with KAPA HiFi and Qiagen Taq, a 5 minute 72**°**C final extension step was used as opposed to 10 minutes. For V4 amplification with KAPA HiFi and Qiagen Taq, no 72**°**C final extension step was used as opposed to 10 minutes.

The following primers were used (16S-specific sequences in bold):

<u>V4 amplicon:</u>

V4_515F_Nextera: TCGTCGGCAGCGTCAGATGTGTATAAGAGACAG**GTGCCAGCMGCCGCGGTAA**

V4_806R_Nextera: GTCTCGTGGGCTCGGAGATGTGTATAAGAGACAG**GGACTACHVGGGTWTCTAAT**

<u>V3V4 amplicon:</u>

V3F_Nextera: TCGTCGGCAGCGTCAGATGTGTATAAGAGACAG**CCTACGGGAGGCAGCAG**

V4_806R_Nextera: GTCTCGTGGGCTCGGAGATGTGTATAAGAGACAG**GGACTACHVGGGTWTCTAAT**

<u>V4V6 amplicon:</u>

V4_515F_Nextera: TCGTCGGCAGCGTCAGATGTGTATAAGAGACAG**GTGCCAGCMGCCGCGGTAA**

V6R_Nextera: GTCTCGTGGGCTCGGAGATGTGTATAAGAGACAG**CGACRRCCATGCANCACCT**

<u>V1V3 amplicon:</u>

V1_27F_Nextera: TCGTCGGCAGCGTCAGATGTGTATAAGAGACAG**AGAGTTTGATCMTGGCTCAG** V3_534R_Nextera: GTCTCGTGGGCTCGGAGATGTGTATAAGAGACAG**ATTACCGCGGCTGCTGG** The resulting amplicons were indexed using KAPA HiFi, normalized, quantified, and sequenced as described above.

### ITS1 primer tests

ITS1 libraries were prepared as previously described (9), using conditions for KAPA HiFi HotStart 0.5x enzyme concentration described above. The following primers were used to compare amplification with (ITS1F*_Nextera/ITS2*_Nextera) and without ITS1F_Nextera/ITS2_Nextera) phosphorothioate protection:

ITS1F_Nextera: TCGTCGGCAGCGTCAGATGTGTATAAGAGACAG**CTTGGTCATTTAGAGGAAGTAA**

ITS2_Nextera: GTCTCGTGGGCTCGGAGATGTGTATAAGAGACAG**GCTGCGTTCTTCATCGATGC**

ITS1F*_Nextera: TCGTCGGCAGCGTCAGATGTGTATAAGAGACAG**CTTGGTCATTTAGAGGAAG*TAA**

ITS2*_Nextera: GTCTCGTGGGCTCGGAGATGTGTATAAGAGACAG**GCTGCGTTCTTCATCGA*TGC**

The resulting amplicons were indexed using KAPA HiFi, 5 µl of the resulting amplicons were run on a 2% agarose gel to examine primer dimer formation.

### Collection and extraction of skin microbiome samples

Skin samples from different body sites (scalp, face, armpit, toe web, and forearm) were collected using either prototype or commercially released OMR-140 OMNIgene•SKIN collection devices, according to manufacturer’s instructions. Samples were treated with 5 µl of Proteinase K (80mg/mL) and incubated at 50°C in a water bath for 1h, then DNA was extracted either using the Qiagen PowerSoil Pro Kit or an optimized bead beating based low biomass extraction method co-developed by DNA Genotek and Diversigen. Briefly, the entire volume (1mL) of an OMR-140 collected sample is transferred to a bead beating containing a mix of 0.5 and 0.1mm glass beads and then homogenized for 10min. Following inhibitor removal, the DNA is purified on a silica column, washed and finally eluted in nuclease free water.

### Preparing 16S rRNA gene variable region V4 or V1V3 sequencing libraries from skin microbiome samples

Skin sample 16S rRNA gene libraries were prepared and sequenced by Diversigen, Inc. Briefly, samples were quantified via qPCR using primers for variable region 4 of the 16S rRNA gene (V4_515F/V4_806R). Samples were amplified as previously described (9) using either KAPA HiFi polymerase or Qiagen Taq and primers for either the V4 (V4_515F/V4_806R) or V1V3 (V1_27F/V3_534R) variable regions. The resulting amplicons were indexed using KAPA HiFi polymerase, normalized, quantified, and sequenced as described above.

### Data Analysis

#### Primer editing analysis

Analysis of edits to primer sequences was carried out as previously described (9). Mismatches to the V4 primer sequences were identified using custom Python scripts and BioPython (17). Illumina adapters were trimmed using cutadapt (18) and paired reads were merged using PANDAseq (19). The first 19 bases (V4_F primer) or 20 bases (V4_R primer) were then compared to the reference sequences and mismatching bases were enumerated. In order to filter out noise from indels in the primer regions, a threshold of a maximum of three mismatches per read was used for this analysis. The proportion of mismatched bases observed at each position was divided by the proportion of expected variants in the primer editing standard plasmid pool as assessed by REcount measurements (16) to generate the Observed edits/Expected edits metric. Sequencing data was alternatively analyzed by counting the number of bases at each position in the primer sequence. The results of these analyses were consistent with the Observed edits/Expected edits metric reported in the paper. Scripts used to analyze the primer editing standards are available on GitHub (https://github.com/darylgohl/PrimerEditing).

#### HM-276D mock community analysis

Fastq files were evenly subsampled down to a maximum of 50,000 reads per sample. Primer and adapter sequences were trimmed off the ends of reads using cutadapt (18). Forward and reverse reads were stitched using PANDAseq (19). The resulting reads were mapped to an HMP mock community reference file (20) using the BURST aligner (version: embalpha_0.99.3_mac) (21).

#### Skin microbiome analysis

Fastq files were evenly subsampled down to a maximum of 50,000 reads per sample. Primer sequences were trimmed off the 5’ ends of reads using cutadapt (18), reads without a primer were discarded (V4 command: cutadapt -g ^GTGCCAGCMGCCGCGGTAA –G ^GGACTACHVGGGTWTCTAAT -e .2 discard-untrimmed; V1V3 command: cutadapt –g ^AGAGTTTGATCMTGGCTCAG -G ^ATTACCGCGGCTGCTGG -e .2 discard-untrimmed). Adapter sequences were trimmed off the 3’ ends of reads using cutadapt (18) (cutadapt -a CTGTCTCTTATACACATCTCCGAGCCCACGAGAC –A CTGTCTCTTATACACATCTGACGCTGCCGACGA). Samples were denoised using DADA2 (22) (qiime dada2 denoise-paired p-trim-left-f 0 p-trim-left-r 0 p-trunc-len-f 167 p-trunc-len-r 167). A phylogenetic tree was created using qiime phylogeny align-to-tree-mafft-fasttree (23). Taxonomy was assigned using qiime feature-classifier classify-sklearn (23).

## RESULTS

### Design and construction of primer editing standard pool

In order to characterize the phenomenon of primer editing in more detail, we designed and constructed a set of 31 primer editing standard plasmids containing the wild type *Escherichia coli* (*E. coli*) V4_515F primer binding site as well as every possible single base mismatch within the last 10 nucleotides of the *E. coli* V4_515F primer binding site (Figure 1A-B). The primer editing standard plasmids also include the *E. coli* 16S rRNA gene variable region 4 sequence and the wild type V4_806R primer binding site. In addition, each of these plasmids contains a unique REcount barcode construct which allows highly-accurate PCR-free measurement of the composition of the standard pool (16). The primer editing plasmids were sequence-verified and pooled evenly. The composition of the primer editing pool was verified by digesting the pool with *Mly*I to liberate the REcount barcode constructs, sequencing, and determining the percent abundance for each plasmid barcode (Figure 1C).

The primer editing standard plasmid pool was then used to create an amplicon sequencing library with KAPA HiFi polymerase, which we previously demonstrated has primer editing activity (9), using primers corresponding to the wild-type *E. coli* V4_515F and V4_806R primer binding sites. In the resulting sequencing reads, substantial editing of the forward amplification primer was observed for the last 7 bases of the primer sequence. Between 8-13% of the sequencing reads contained a non-wild-type base with the last 7 bases of the amplification primer (Figure 1C), consistent with the composition of the template sequences where variants were present at each position in 3/31 plasmids (9.68%). The percentage of edits observed within the last 6 bases of the primer sequence tracked the REcount abundance measurements, suggesting that editing at these positions was near complete, while the 7th base from the 3’ end was likely incompletely edited. Primer editing was barely detected 8 bases from the 3’ end of the primer. As expected, no appreciable primer editing was observed for the wild type reverse primer reads, in which the template sequence perfectly matched the amplification primer in all of the primer editing standard plasmids (Figure 1D).

### Ability of different proofreading polymerases to mediate primer editing

In previous work, we demonstrated that both KAPA HiFi and NEB Q5 polymerases exhibit primer editing activity (9). We used the primer editing standards to explore whether other commercially available enzymes were able to mediate primer editing (Figure 2A). In addition to KAPA HiFi and Q5, we tested three additional proofreading enzymes (Pfu, Phusion, and Vent polymerase) and three non-proofreading polymerases (NEB Taq, Qiagen Taq, and Accuprime Taq). All five proofreading polymerases exhibited some level of primer editing activity, though the extent of editing observed varied for each proofreading polymerase, ranging from near complete editing of up to 6 bases for KAPA HiFi polymerase to near complete editing of only the last 2 bases for Phusion polymerase (Figure 2A). No appreciable primer editing activity was observed for any of the non-proofreading Taq polymerases tested (Figure 2A).

**Figure 2.**
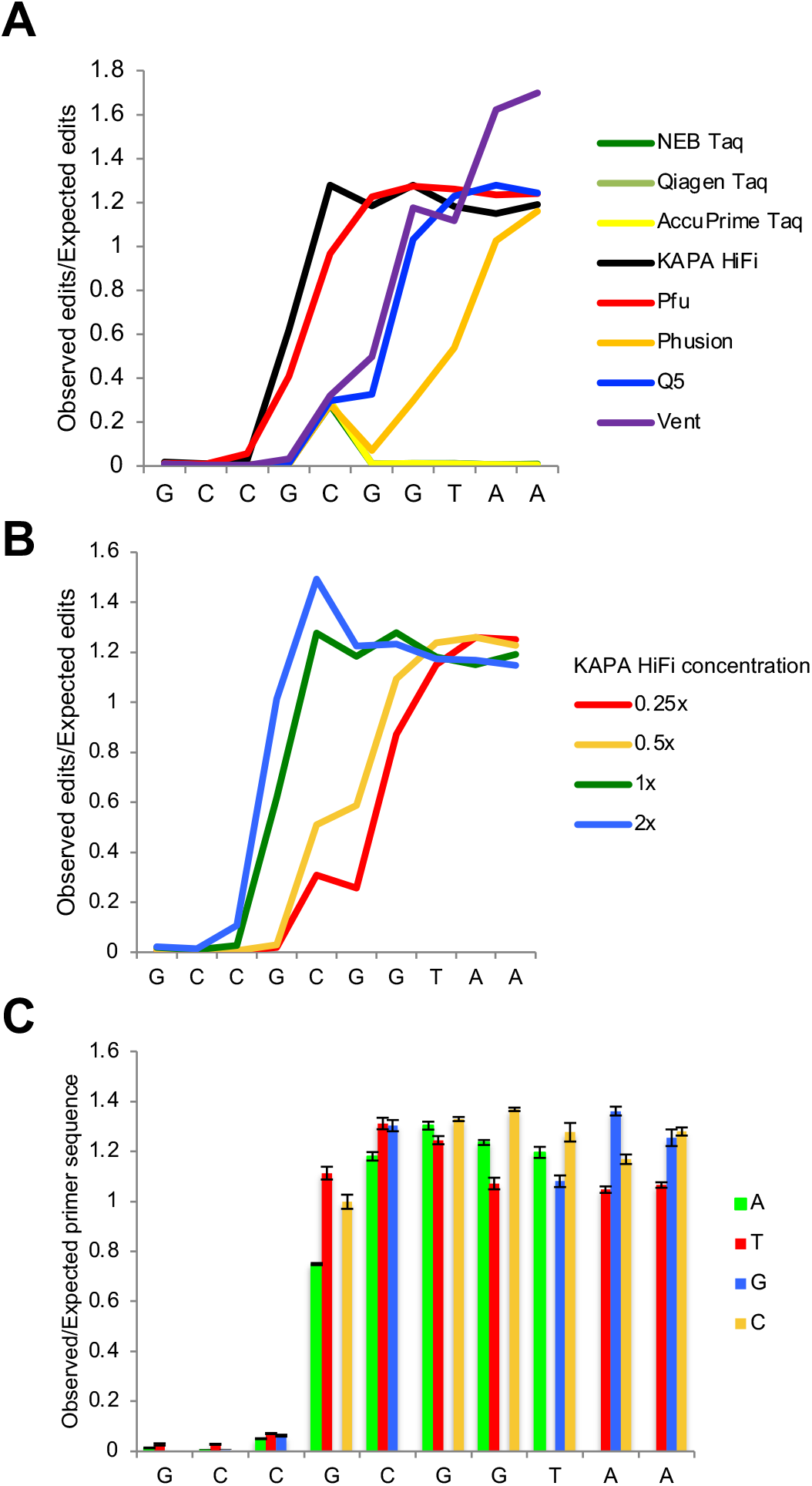
Assessing the effect of different polymerases, polymerase concentration, and sequence identity on primer editing. A) Ratio of observed versus expected edits position in the forward primer sequence when amplified with an *E. coli*-specific 515F/806R primer set and the indicated polymerase. B) Ratio of observed versus expected edits position in the forward primer sequence when amplified with an *E. coli*-specific 515F/806R primer set and KAPA HiFi polymerase at the indicated enzyme concentration. C) Ratio of observed versus expected edits for each individual base of the forward primer sequence when amplified with an *E. coli*-specific 515F/806R primer set and 1x KAPA HiFi polymerase (n=3, error bars = +/-S.E.M.).

### Primer editing is dependent on enzyme concentration

We previously hypothesized that the primer editing activity of KAPA HiFi polymerase was dependent on enzyme concentration (9). We generated sequencing libraries by amplifying the primer editing standard plasmid pool with each of the enzymes tested above across a range of enzyme concentrations (0.25x, 0.5x, 1x, and 2x the manufacturer’s recommended concentration). Consistent with our previous observations, the extent of primer editing exhibited a dependence on the concentration of the KAPA HiFi polymerase in the reaction (Figure 2B). A similar concentration dependence was observed with the other four proofreading polymerases tested (Supplemental Figure S1).

### Primer editing exhibits minimal sequence-specificity

Minimal sequence specificity was observed in primer editing (Figure 2C). In general, there was no clear pattern in the base composition of the primer edits, with the possible exception that a slightly increased proportion of G and C edits were observed at the 3’ end of the primer, relative to T edits (Figure 2C). It is possible that this reflects a small increase in amplification efficiency due to the presence of a stronger G:C bond in the 3’ end of the primer.

### Effect of primer mismatches on amplification efficiency

Next, we examined the extent to which primer editing is expected to rescue the amplification of templates that have primer mismatches. Mismatches within the last 3 bases of the primer sequence have been shown to be the most deleterious to amplification efficiency (10, 11). To examine the effect of the individual primer mismatches on amplification efficiency, we used a pool of primers that contained the same 31 variants as the primer editing standard plasmids (variant primers) to amplify an *E. coli* template with wild type V4_515F and V4_806R primer binding sites with either KAPA HiFi polymerase or a non-proofreading Taq polymerase. For KAPA HiFi polymerase, we observed edits of the variant primers to match the wild type template sequence in a pattern that was roughly complementary to the editing of the wild type primer to match the variants encoded by the primer editing standard plasmids (Figure 3A). For the non-proofreading Taq polymerase, where no editing of the wild-type primer to match the primer editing standards is seen, we examined the proportion of variant primers able to amplify the wild-type template as a measure of the amplification penalty incurred by each mismatch (Figure 3B). Variant primers at all positions were permissive for amplification by Taq at some level. However, variants within the last five bases of the forward amplification primer exhibited notable amplification penalties as reads containing these variants were present at 0.922, 0.774, 0.404, 0.318, 0.189-fold their expected values. We also tested whether primer editing or the permissiveness of amplification of primer mismatches were sensitive to annealing temperature by performing the above amplifications across a gradient of annealing temperatures ranging from 50°C to 60°C. Within this range of annealing temperatures, there were no effects on the amount of primer editing or the amount of amplification by variant primers (Supplemental Figure S2). Thus, primer editing helps to overcome reduced amplification efficiencies due to primer mismatches within the last 4-5 bases of the amplification primer, while in the case of the V4_515F primer variants tested, variants beyond 5 bases from the 3’ end of the primer were generally permissive for amplification.

**Figure 3.**
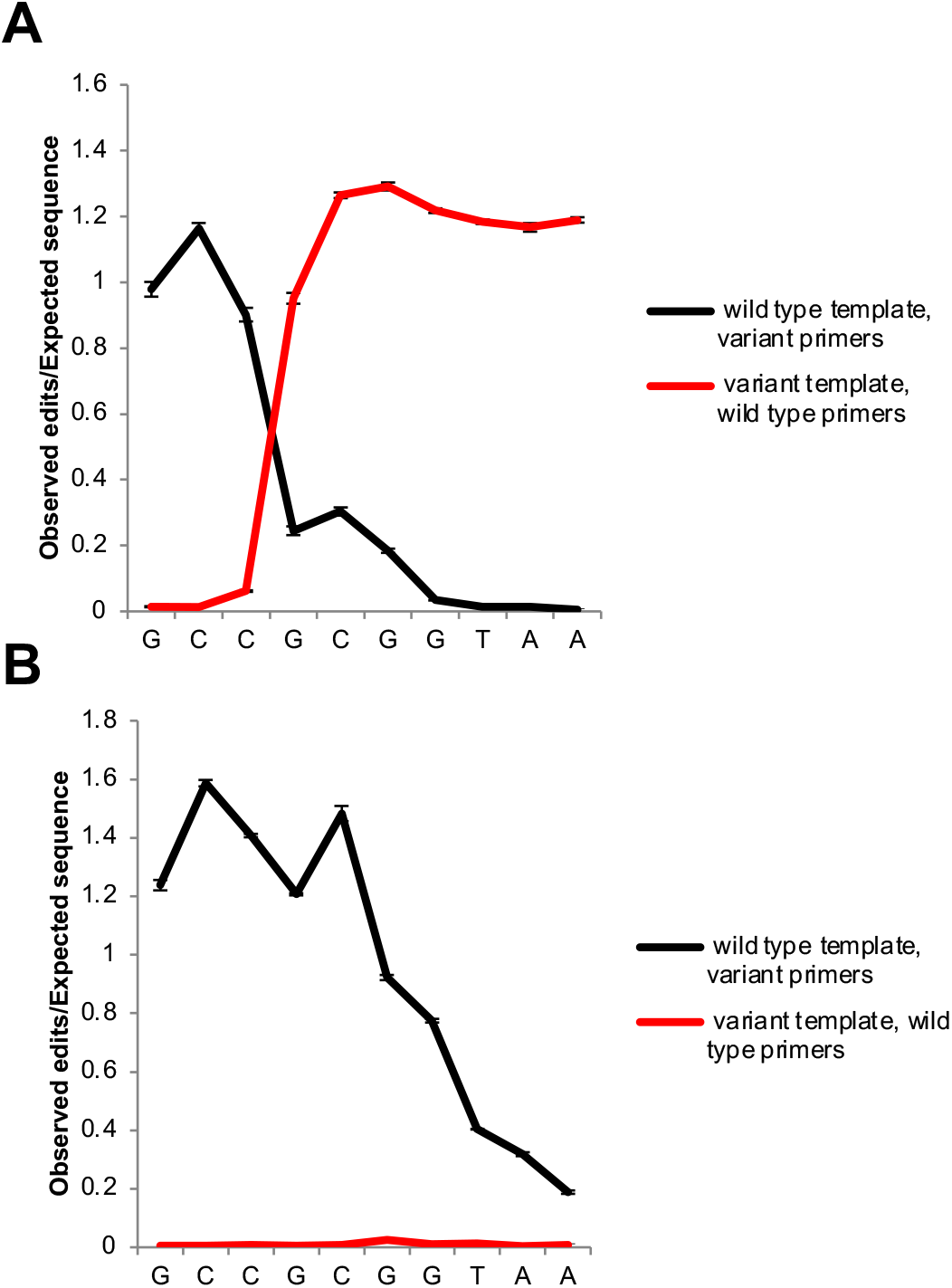
Reciprocal editing of mutant and wild type primers. Comparison of edited wild type *E. coli*-specific 515F primers used to amplify the primer editing standards and a collection of mutant primers which mirror the composition of the primer editing standards used to amplify an *E. coli* wild type template using either A) KAPA HiFi polymerase or B) Taq polymerase. (n=3, error bars = +/-S.E.M.).

### Primer editing is tunable through the incorporation of phosphorothioate bonds in amplification primers

Phosophorothioate bonds (in which a non-bridging oxygen within the oligonucleotide phosphate backbone is replaced with a sulfur) have been previously shown to block the activity of various exonucleases and are commonly used to prevent exonuclease degradation of oligonucleotides (24–26). Using the primer editing standard plasmid pool, we tested whether phosphorothioate bonds could block primer editing by preventing the 3’ to 5’ exonuclease activity of proofreading polymerases. For KAPA HiFi polymerase, Q5 polymerase, and Phusion polymerase, a single phosphorothioate bond was able to block primer editing beyond the position of the phosphorothioate bond (Figure 4, Supplemental Figure S3). Thus, the extent of primer editing can be tuned through the strategic placement of phosphorothioate bonds in the amplification primers. This ability to limit the extent of primer editing can be used to prevent the formation of undesirable reaction products such as primer dimers. We tested the incorporation of phosphorothioate bonds into an ITS1 (Internal Transcribed Spacer) primer set used for fungal microbiome profiling where we previously saw extensive primer dimer formation during amplification with KAPA HiFi polymerase. The addition of phosphorothioate bonds between the third and fourth to last bases of the forward and reverse primers eliminated the formation of primer dimers for this ITS1 primer set (Figure 5).

**Figure 4.**
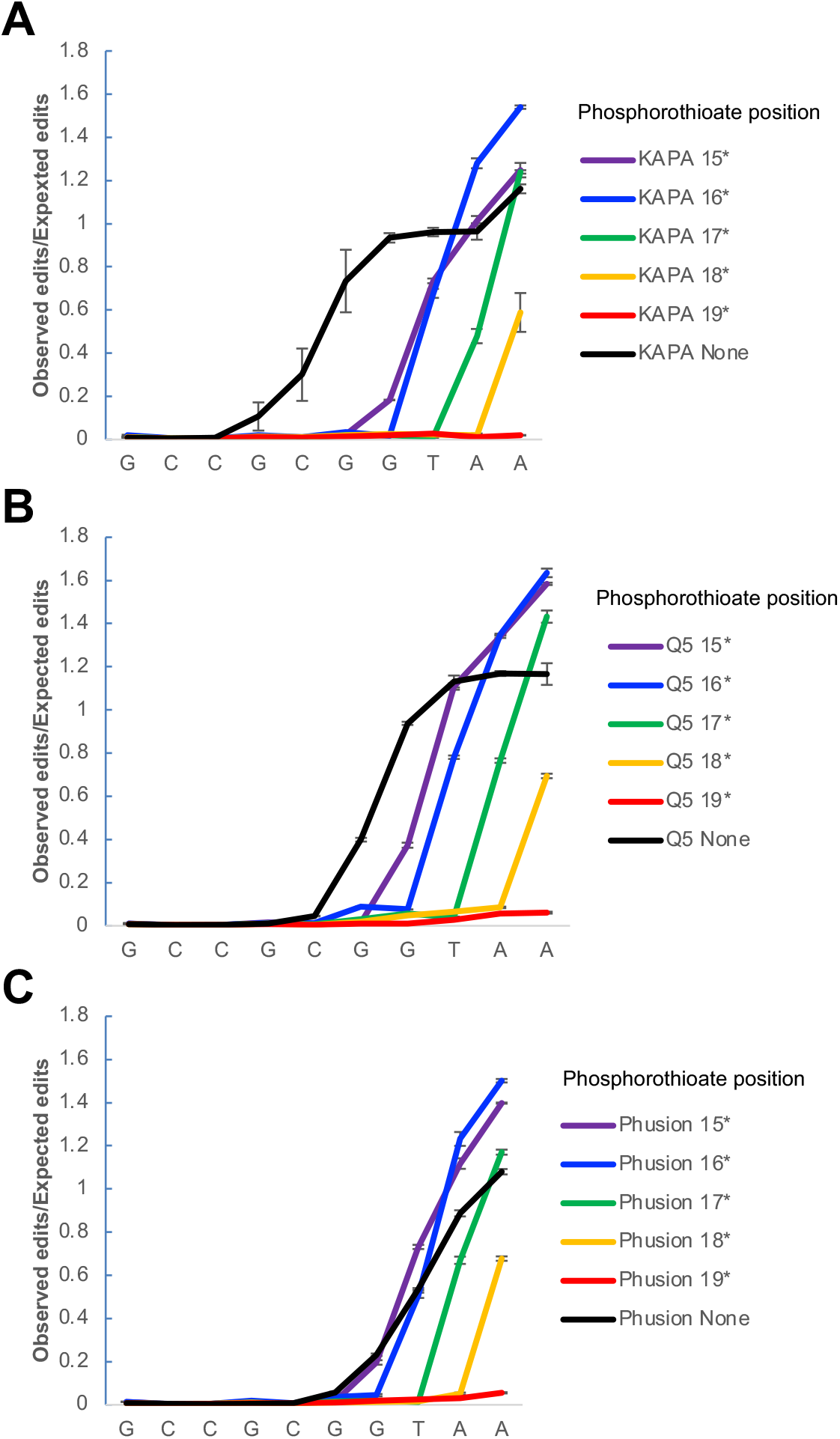
Tuning of primer editing using phosphorothioate protection. Effect of incorporating phosphorothioate bonds into *E. coli*-specific 515F primers on extent of primer editing observed when the primer editing standards are amplified using A) KAPA HiFi polymerase (n=3, error bars = +/-S.E.M.); B) NEB Q5 polymerase (n=3, error bars = +/-S.E.M.); C) Phusion polymerase (n=3, error bars = +/-S.E.M.).

**Figure 5.**
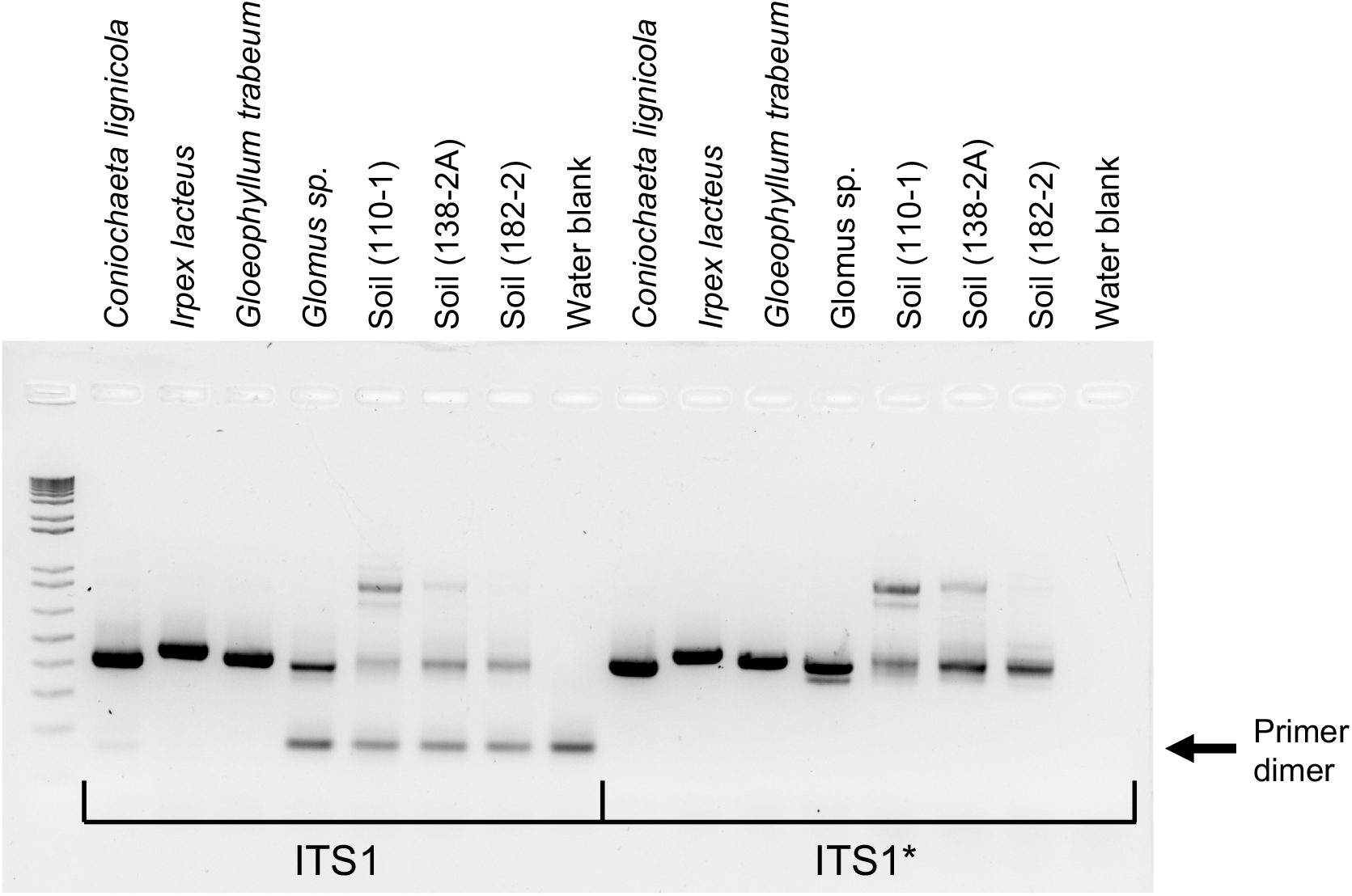
Improving performance of ITS1 primers. Fungal isolates and soil samples amplified with unprotected (left, ITS1) or phosphorothioate protected (right, ITS*) primers. Primer dimer bands are indicated by the arrow.

### Effect of primer mismatches on amplification of DNA mock community samples

Using DNA-based mock microbial communities and other non-human primate and human samples, we previously demonstrated that primer editing could mitigate the drop-out of taxa due to primer mismatches in amplicon-based microbiome profiling (9). We amplified a mock microbial community made by Human Microbiome Project (HM-276D) using each of the eight polymerases tested in Figure 2. As expected, the three non-proofreading polymerases (AccuPrime Taq, NEB Taq, Qiagen Taq) were unable to amplify *Cutibacterium acnes* (*C. acnes*), which has mismatches between the 16S rRNA gene template the 3’ end of the V4 variable region amplification primers (Figure 6A). All five proofreading enzymes were able to amplify *C. acnes*, despite the primer mismatches; the recovery of *C. acnes* was reduced for the sample amplified with Phusion polymerase (Figure 6A), consistent with the weaker primer editing activity observed with this enzyme with the synthetic primer editing standards (see Figure 2A).

**Figure 6.**
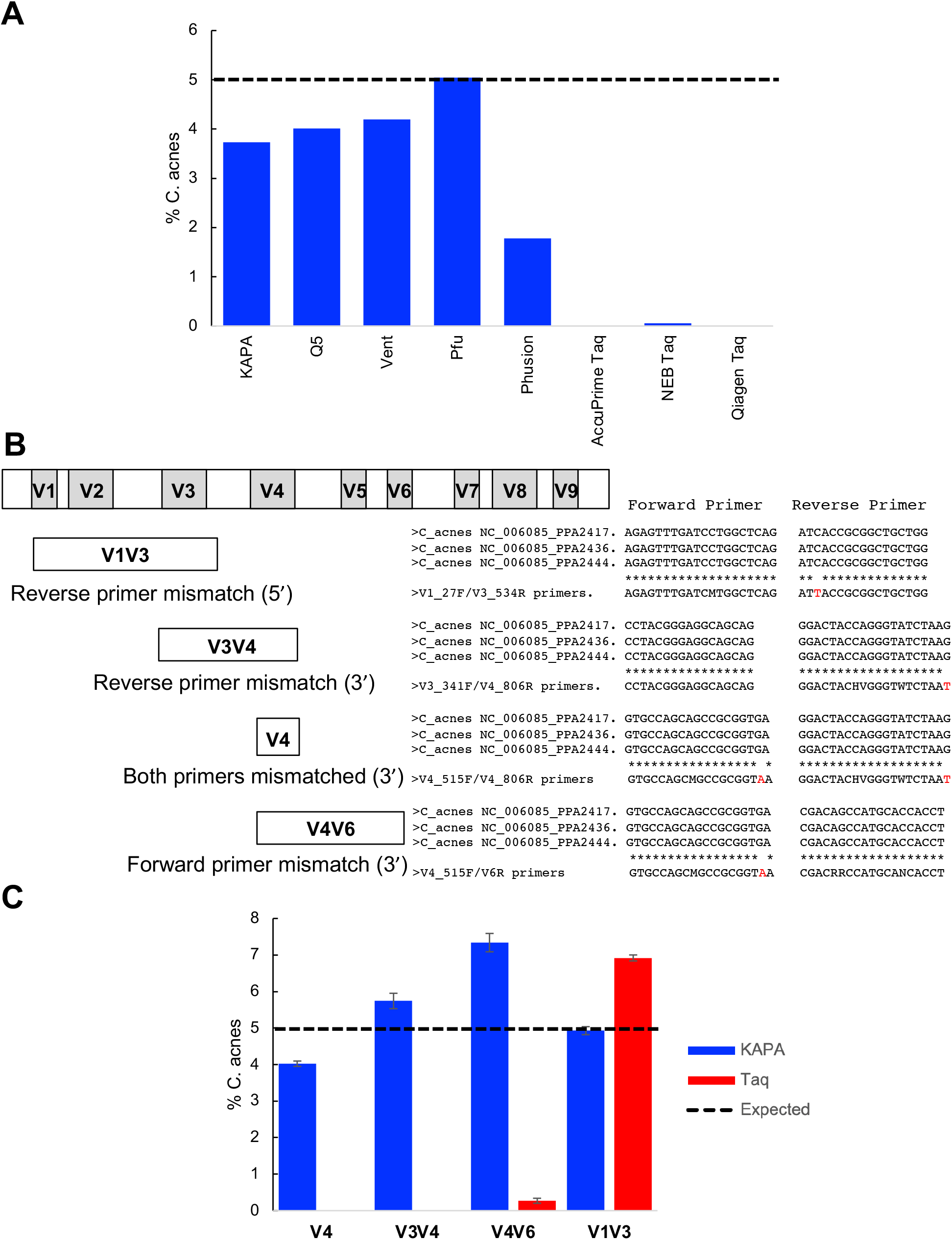
Primer editing recovers *C. acnes* across multiple variable regions. A) Percent *C. acnes* observed in the HM-276D mock community when amplifying the V4 variable region using the indicated polymerase. Expected abundance of *C. acnes* is indicated by the dashed line. B) Schematic of the 16S rRNA gene with variable regions indicated. The status of primer mismatches to the *C. acnes* template sequence for the primers used to amplify the V1V3, V4, V3V4, and V4V6 variable regions is indicated. C) Percent *C. acnes* observed in the HM-276D mock community when amplifying different variable regions using either KAPA HiFi polymerase or Qiagen Taq polymerase (n=3, error bars = +/-S.E.M.). Expected abundance of *C. acnes* is indicated by the dashed line.

Other 16S rRNA gene variable regions besides the V4 variable region are often used for specific sample types, as different research communities strive to optimize the taxonomic resolution or minimize the drop-out of key taxa. For instance, the V1V3 variable regions have been recommended for the profiling of human skin associated communities on the basis of improved resolution of key taxa and due to the fact that drop-out of *C. acnes* occurs when the V4 variable region is used in the absence of primer editing (27, 28). We examined the effect of using primers targeting different variable regions on the amplification of the Human Microbiome Project mock community (HM-276D) (Figure 6B). The detection of near-expected levels of *C. acnes* was dependent on using a proofreading polymerase for primer sets targeting the V4 variable region and also the V3V4 and V4V6 variable regions, both of which contain 3’ mismatches in one of the two amplification primers (Figure 6C, Supplemental Figure S4). As previously reported (27), *C. acnes* template was amplified by V1V3 primers, both in the presence or absence of primer editing (Figure 6C, Supplemental Figure S4). The V1V3 reverse primer contains a single mismatch to the *C. acnes* template near the 5’ end primer which would not be expected to interfere with primer annealing or extension in the absence of primer editing (see Figure 3B).

### Primer editing enables robust detection of *Cutibacterium* in skin microbiome samples with the 16S rRNA gene V4 variable region

In order to determine whether primer editing by a proofreading polymerase can overcome the drop-out of *C. acnes* observed in human skin microbiome samples with primers targeting the 16S rRNA gene V4 region, we compared the V4 and V1V3 microbiome profiles from a number of skin body sites amplified using either the proofreading KAPA HiFi polymerase or the non-proofreading Taq polymerase (Figure 7). As expected (27), substantial levels of the *Cutibacterium* genus was observed for a variety of skin body sites, including face (Figure 7A), forearm (Figure 7B), armpit (Figure 7C), and scalp (Figure 7D) when these samples were amplified with the V1V3 primers. This was true for amplification using both a proofreading polymerase (KAPA HiFi) and a non-proofreading polymerase (Qiagen Taq). A greater proportion of *Cutibacterium* was observed in the case of amplification of the V1V3 region with Taq polymerase (Supplemental Figure S5) and for these samples, it is not certain which enzyme is more reflective of the actual composition of the samples. However, a similar effect was seen for the HMP mock community, where the amount of *C. acnes* observed when the V1V3 variable region was amplified with Taq polymerase was 40% higher than when amplified KAPA HiFi (Figure 6B). In the case of the HMP mock community, the KAPA HiFi measurements were closer to the expected 5% abundance of *C. acnes*, suggesting that for the V1V3 amplicon the abundance of *Cutibacterium* may be overestimated when amplified with Taq polymerase (Supplemental Figure S5).

**Figure 7.**
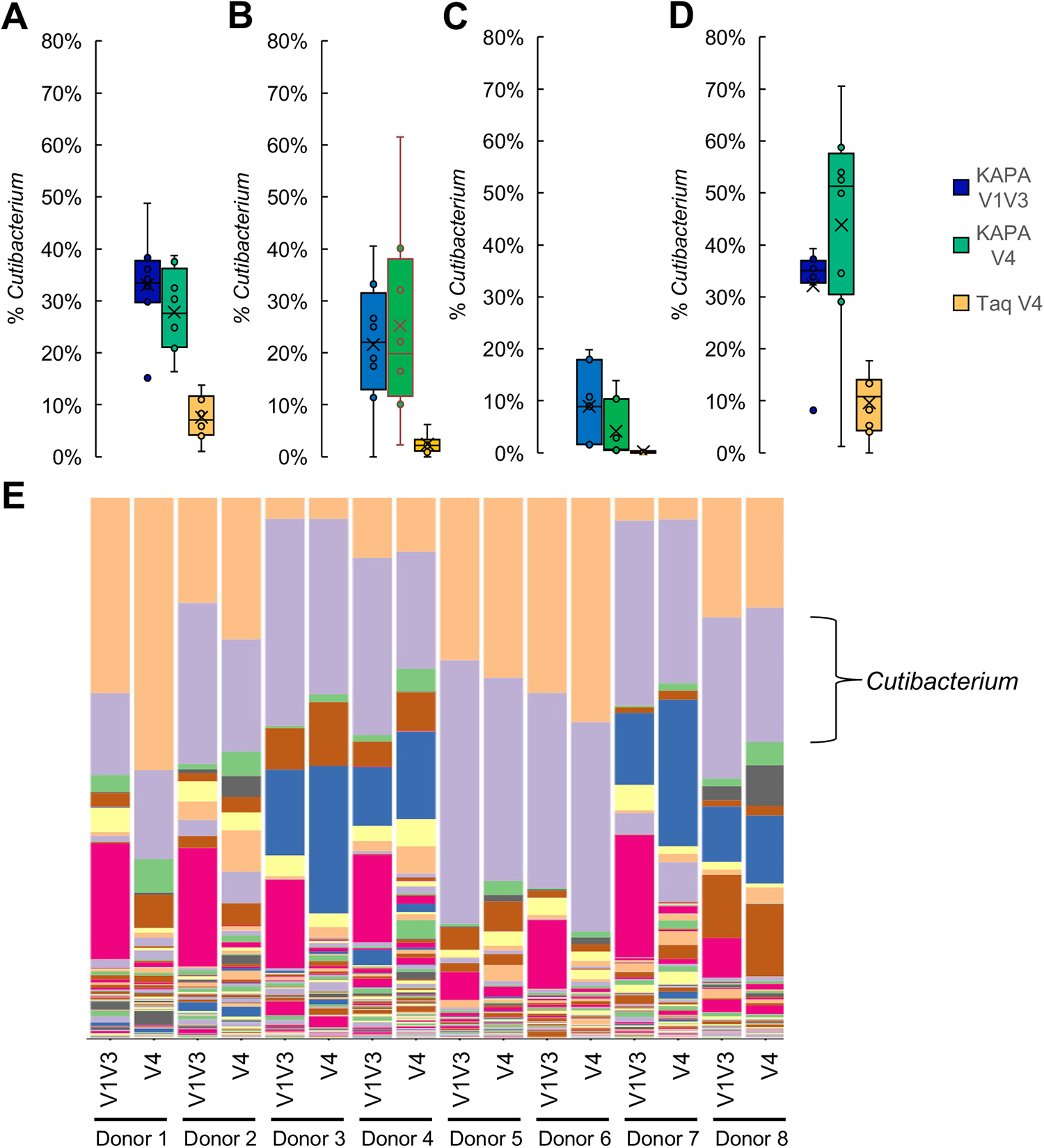
Recovery of *C. acnes* in skin microbiome samples using primer editing. A) Percent *Cutibacterium* observed in face microbiome samples (n=8) amplified with the indicated conditions. B) Percent *Cutibacterium* observed in forearm microbiome samples (n=8) amplified with the indicated conditions. C) Percent *Cutibacterium* observed in armpit microbiome samples (n=7) amplified with the indicated conditions. D) Percent *Cutibacterium* observed in scalp microbiome samples (n=8) amplified with the indicated conditions. E) Microbiome profiles for face samples from 8 donors amplified with either V4 or V1V3 primers using KAPA HiFi polymerase. *Cutibacterium* genus is shown in purple. For full legend and visualization of all skin microbiome data see Supplemental File 4. For box plots, mean is indicated by “X”, median is indicated by horizontal line, data points are indicated by dots, boxes span the second and third quartiles, and the data range (min and max) is indicated by the whiskers or outlier data points.

When this set of skin microbiome samples was amplified using the 16S rRNA gene V4 primers, the abundance of *Cutibacterium* was substantially decreased for samples amplified with Taq polymerase, likely due to the reduced amplification efficiency caused by the mismatches between the V4 primers and the *Cutibacterium* 16S template (Figure 7A-D). In contrast, when the same skin samples were amplified using the 16S rRNA gene V4 primers and KAPA HiFi polymerase, similar levels of *Cutibacterium* were observed as when the V1V3 region was amplified with KAPA HiFi (Figure 7A-E). Thus, the incorporation of primer editing into the design of amplicon-based microbiome experiments can mitigate the effect of taxa dropout due to primer mismatches and in the case of human skin microbiome samples enables the recovery of similar levels of *C. acnes* using 16S rRNA gene V4 and V1V3 primers.

## DISCUSSION

Here we report a novel set of synthetic standards that allowed us to examine the phenomenon of primer editing by proofreading polymerases in detail. We use these synthetic standards to demonstrate that a variety of proofreading polymerases, including KAPA HiFi, NEB Q5, Phusion, Pfu, and Vent, can all mediate primer editing to enable efficient amplification of templates which have mismatches to the amplification primers (Figure 2). The amount of primer editing observed varies between the different enzymes, is sensitive to enzyme concentration, and exhibits minimal sequence specificity (Figure 2).

Kinetic studies of proofreading polymerases such as the T7 DNA polymerase have demonstrated that selective exonucleolytic repair of mismatches relies on kinetic partitioning between the polymerase and exonuclease sites (29) where incorporation of a mismatch slows the rate of polymerization (30), allowing transfer of the mismatched DNA to the exonuclease site from the normally thermodynamically favored polymerase site (29, 31). Modern engineered polymerases such as KAPA HiFi and NEB Q5 polymerase are the result of targeted protein engineering and directed evolution (32–34). During these processes, enzymes are selected on the basis of a number of desirable properties, such as error rate, GC bias, and processivity. It is possible that the kinetic parameters that determine the balance of exonuclease and polymerase activity were intentionally or inadvertently altered during this selection process. While determining the detailed mechanisms of the differences in primer editing exhibited by different enzymes will be an interesting topic for future studies, we speculate that enzymes such as KAPA HiFi may have increased exonuclease activity or accessibility that explain its enhanced primer editing activity and may also correlate with other desirable enzyme properties (such as lower error rates).

Using the synthetic primer editing standards, we determined that for some enzymes (KAPA HiFi and Pfu) near-complete editing of mismatches could extend as far as 6 bases from the end of the primer (Figures 1-2). Kinetic studies have demonstrated that mismatches or lesions in the n-1 to n-4 positions can induce polymerase stalling (35–37) and structural studies using the proofreading *Bacillus stearothermophilus* DNA polymerase large fragment indicated that mismatches as far as 4 bases from the point of incorporation can disrupt the polymerase active site, leading to a “short-term memory” of mis-incorporated bases (38). To enable primer editing at the n-6 position, it is possible that enzymes such as KAPA HiFi and Pfu have active sites that are sensitive to perturbation by mismatches over longer ranges. Alternatively, non-specific degradation or excision at the 3’ end of the primer could help explain the detection and editing of n-6 mismatches and is also consistent with the elevated levels of primer dimer formation often seen with proofreading enzymes.

The extent of primer editing is tunable through the incorporation of phosphorothioate bonds in the amplification primers, which block the polymerase 3’ to 5’ exonuclease activity (Figure 4). Thus, the primer editing standard plasmid pool enabled us to carry out DNA sequencing-based enzymology and further characterize and refine the utility of primer editing. The incorporation of strategically placed phosphorothioate bonds allows one to take advantage of the benefits of primer editing while reducing detrimental side-effects such as formation of primer dimers (Figure 5), or degradation of primer specificity.

The introduction of a phosphorthioate bond creates a chiral center in the oligonucleotide backbone and during phosphoramidite synthesis roughly equal proportions of Rp and Sp diastereomers are incorporated (24). Many nucleases are sensitive to the chirality of the phosphorothioate bond and multiple consecutive phosphorothioate bonds are often used to completely block exonuclease activity (25, 26). The fact that a single phosphorothioate bond was able to block essentially all primer editing suggests that the 3’ to 5’ exonuclease activity of the proofreading polymerases we tested is either not sensitive to the chirality of the phosphorothioate bond or that the cumulative effect of random incorporation of the blocking stereoisomer over the course of multiple PCR cycles is able to substantially inhibit the efficiency of amplifying edited primers beyond the site of phosphorothioate incorporation.

Using both mock microbial communities and human skin microbiome samples we show that primer editing can be used to minimize the dropout or under-amplification of taxa with mismatches to amplification primers across a number of commonly used microbiome primer sets. In particular, we demonstrated that primer editing can overcome previously documented shortcoming of the V4_515F/V4_806R primer set in amplifying *C. acnes* from skin microbiome samples (27, 28), recovering similar levels of *Cutibacterium* as those observed with V1V3 primers (Figure 7). It should be noted that suitable sequencing strategies are also required to successfully employ primer editing in microbiome experiments, as we previously demonstrated that using custom sequencing primers can negate the beneficial effects of primer editing (9). *In silico* analysis and comparison to shotgun microbiome data suggests that primer mismatches in the critical final 3-4 bases of commonly used amplification 16S rRNA gene amplification primers are relatively common (12), and the discovery of new phyla with divergent 16S rRNA gene sequences (15) further support the use of amplification strategies that can overcome primer mismatches as a means to improve the accuracy of amplicon-based microbiome measurements.

## DATA AVAILABILITY

Sequencing data files are available through the NCBI Sequence Read Archive, BioProject: PRJNA673973.

## ACKNOWLEDGEMENTS

We thank the staff of the University of Minnesota Genomics Center for advice, technical support, and data generation. We thank Megan Johnson, Shea Anderson, and Andrew Cross for help with experiments, Alex LaReau for help with analysis, Sarah Castle and Linda Kinkel for providing samples for the ITS1 primer tests, and Lisa Gamwell for helpful comments. The authors acknowledge the Minnesota Supercomputing Institute (MSI) at the University of Minnesota for providing resources that contributed to the research results reported within this paper.

