## Supplemental Figures for "Dissecting and Tuning Primer Editing by Proofreading Polymerases"

### Supplemental Files

**Supplemental File 1. Design of primer editing constructs**
UMGC_257_Primer_editing_standards_direct_synthesis.xlsx

**Supplemental File 2. REcount barcode mapping file**
UMGC_257_STD_pool_quant_barcodes.fasta

**Supplemental File 3. Variant-containing primers**
UMGC_257_variant_primers.xlsx

**Supplemental File 4. QIIME visualization file for skin microbiome data**
taxa-bar-plots-with-metadata_V1V3&V4.qzv

### Supplemental Figure Legends

**Supplemental Figure S1. Concentration dependence of primer editing.**
Ratio of observed versus expected edits position in the forward primer sequence when amplified with an *E. coli*-specific 515F/806R primer set and A) Pfu polymerase, B) Q5 polymerase, C) Phusion polymerase, or D) Vent polymerase at the indicated enzyme concentration.

**Supplemental Figure S2. Annealing temperature has minimal impact on primer editing.**
A) Comparison of edited wild type *E. coli*-specific 515F primers used to amplify the primer editing standards amplified at different annealing temperatures using KAPA HiFi polymerase (n=3, error bars = +/- S.E.M.).
B) Comparison of a collection of mutant primers which mirror the composition of the primer editing standards used to amplify an *E. coli* wild type template at different annealing temperatures using KAPA HiFi polymerase (n=3, error bars = +/- S.E.M.).
C) Comparison of edited wild type *E. coli*-specific 515F primers used to amplify the primer editing standards amplified at different annealing temperatures using Taq polymerase (n=3, error bars = +/- S.E.M.).
D) Comparison of a collection of mutant primers which mirror the composition of the primer editing standards used to amplify an *E. coli* wild type template at different annealing temperatures using Taq polymerase (n=3, error bars = +/- S.E.M.).

**Supplemental Figure S3. Tuning of primer editing using phosphorothioate protection.**
Repeat experiment for data in Figure 4A showing effect of incorporating phosphorothioate bonds into *E. coli*-specific 515F primers on extent of primer editing observed when the primer editing standards are amplified using KAPA HiFi polymerase. (n=3, error bars = +/- S.E.M.).

**Supplemental Figure S4. Mock community amplification data**
Comparison of amplification of the HM-276D mock community using either Qiagen Taq or Kapa HiFi for the following variable regions: A) V4; B) V3V4; C) V4V6; D) V1V3. Plots show average values for three replicate amplifications. Arrows indicate *C. acnes*.

**Supplemental Figure S5. *Cutibacterium* detected in skin microbiome samples amplified with Taq polymerase.**
A) Percent *Cutibacterium* observed in face microbiome samples (n=8) amplified with the indicated conditions. B) Percent *Cutibacterium* observed in forearm microbiome samples (n=8) amplified with the indicated conditions. C) Percent *Cutibacterium* observed in armpit microbiome samples (n=8) amplified with the indicated conditions. D) Percent *Cutibacterium* observed in scalp microbiome samples (n=8) amplified with the indicated conditions. For box plots, mean is indicated by "X", median is indicated by horizontal line, data points are indicated by dots, boxes span the second and third quartiles, and the data range (min and max) is indicated by the whiskers or outlier data points.

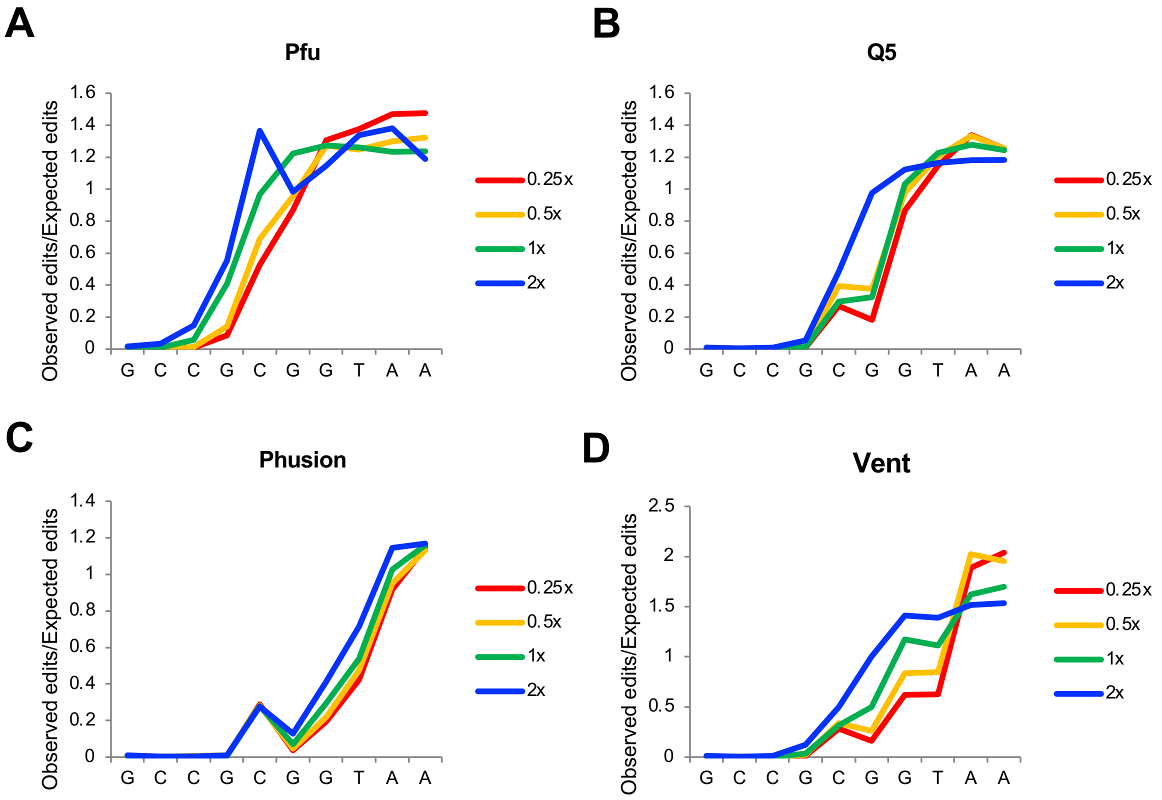


Supplemental Figure S1

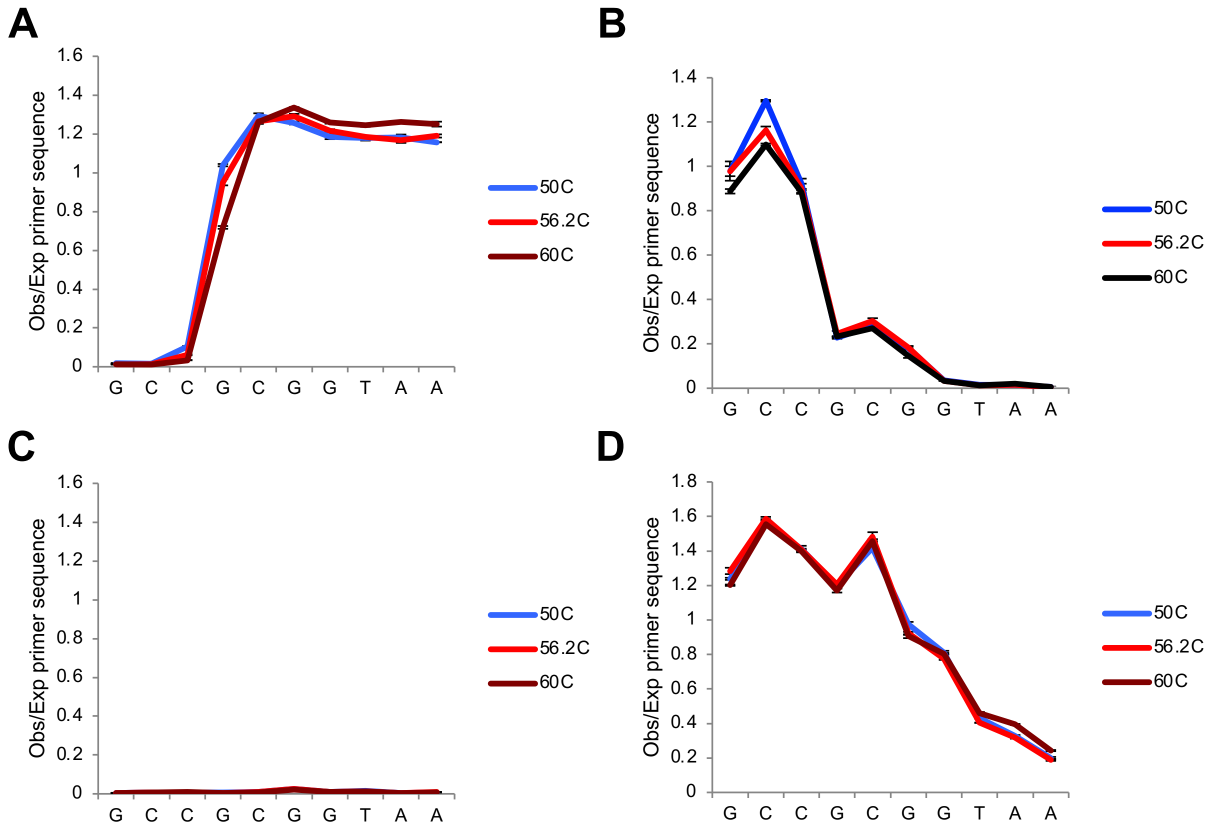


Supplemental Figure S2

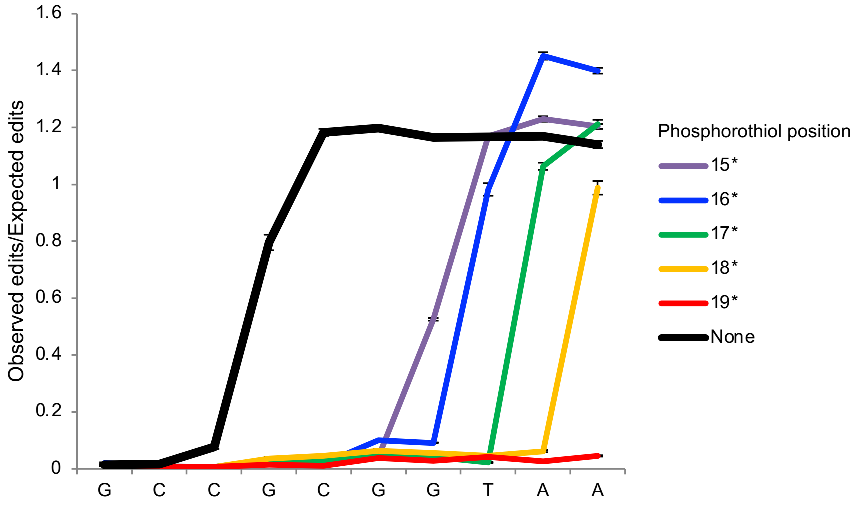


Supplemental Figure S3

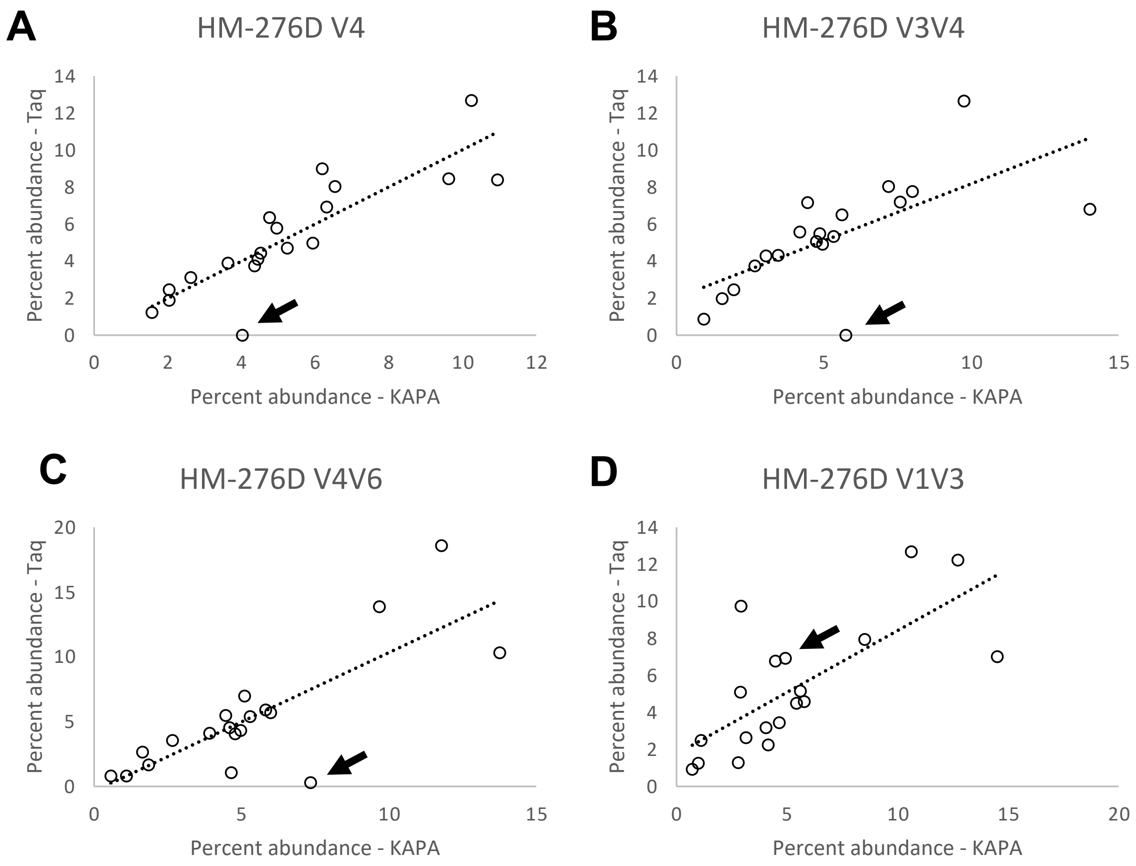


Supplemental Figure S4

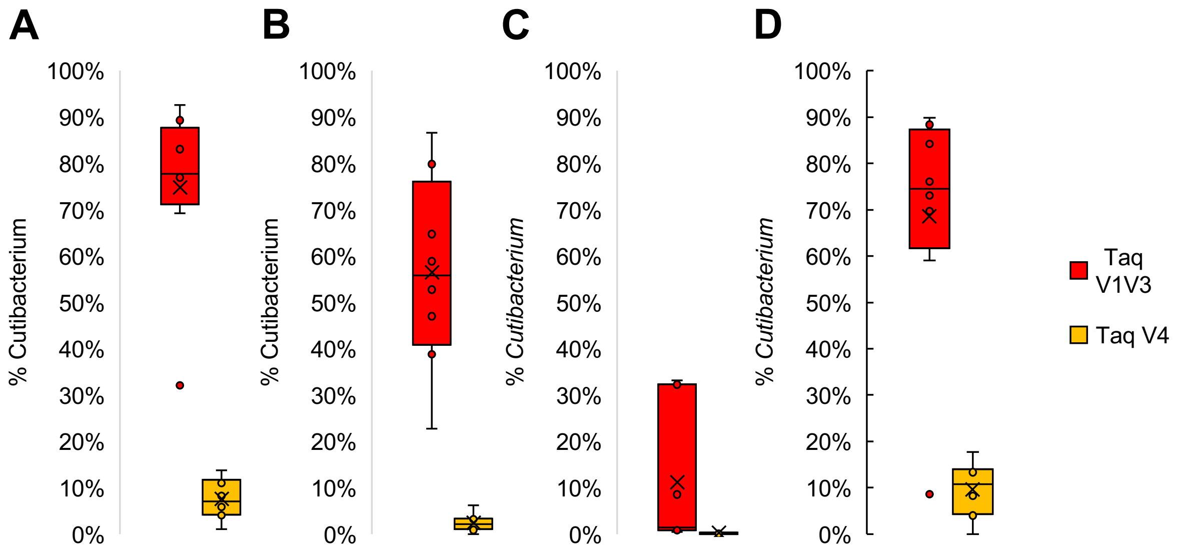


Supplemental Figure S5
